# Aberrant PARP1 Activity Couples DNA Breaks to Deregulated Presynaptic Calcium Signalling and Lethal Seizures

**DOI:** 10.1101/431916

**Authors:** Emilia Komulainen, Jack Badman, Stephanie Rey, Stuart Rulten, Limei Ju, Kate Fennell, Peter J. McKinnon, Kevin Staras, Keith W Caldecott

## Abstract

Defects in DNA single-strand break repair result in cerebellar ataxia which in *Xrcc1^Nes-Cre^* mice is promoted by hyperactivity of the DNA strand break sensor protein, Parp1. Here, we show that Parp1 hyperactivity extends beyond the cerebellum in Xrcc1-defective brain, resulting in lethal seizures and shortened lifespan. We demonstrate that aberrant Parp1 activation triggers seizure-like activity in Xrcc1-defective hippocampus *ex vivo* and aberrant presynaptic calcium signalling in isolated hippocampal neurons *in vitro.* Moreover, we show that these defects are prevented by Parp1 inhibition and/or deletion. Collectively, these data identify aberrant Parp1 activity at unrepaired DNA breaks as a cell-autonomous source of deregulated presynaptic calcium signalling, and highlight PARP inhibition as a possible therapeutic approach in *XRCC1*-mutated neurodegenerative disease.

**Summary:** PARP1 activity and presynaptic Ca^2+^ signalling

## Main text

DNA single-strand breaks (SSBs) are the commonest lesions arising in cells and are rapidly detected by poly(ADP-ribose) polymerase-1 (PARP1) and/or poly(ADP-ribose) polymerase-2 (PARP2); enzymes that are activated at DNA breaks and modify themselves and other proteins with mono-ADP-ribose and/or poly-ADP-ribose *(1–4)*. Poly(ADP-ribose) triggers recruitment of the DNA single-strand break repair (SSBR) scaffold protein XRCC1 and its enzymatic protein partners to facilitate SSBR *(4–8)*. If not repaired rapidly, SSBs can result in replication fork stalling and/or collapse and can block the progression of RNA polymerases during gene transcription *(9–13)*. Moreover, mutations in proteins involved in SSBR are associated in humans with cerebellar ataxia, neurodevelopmental defects, and episodic seizures *(14,15)*. To date, all of the identified SSBR-defective human diseases are mutated in either XRCC1 or in one of its protein partners *(5)*. Recently, we demonstrated that the loss of SSBR in *Xrcc1^Nes-Cre^* mice results in the hyperactivation of PARP1, elevated steadystate levels of poly(ADP-ribose), and cerebellar ataxia *(16)*. However, the mechanism and extent to which PARP1 hyperactivation impacts on neurological function in response to unrepaired endogenous SSBs is unknown.

To examine the extent of Parp1 hyperactivity in Xrcc1-defective brain we measured the steady-state level of ADP-ribose by immunohistochemistry. We detected increased levels of ADP-ribose throughout *Xrcc1^Nes-Cre^* brain, with particularly strong anti-ADP-ribose immunostaining in the cerebellum and hippocampus (Fig.1A & 1B). The elevated ADP-ribose was the product of Parp1 activity because *Parp1* deletion in *Xrcc1^Nes-Cre^* mice reduced this signal to levels similar to or below those in wild-type brain (Fig.1A; *Parp1^-/-^/Xrcc1^Nes-Cre^*). We did not detect elevated ADP-ribose in brain from *Ku70^-/-^* mice, however, in which the primary if not only pathway for DNA double-strand break (DSB) repair in postmitotic cells is defective, confirming that the elevated ADP-ribose in *Xrcc1^Nes-cre^* brain was the result of unrepaired SSBs, not DSBs (Fig.1A). Notably, as observed previously *(17),* Parp1 protein was significantly downregulated in *Xrcc1^Nes-cre^* brain (Fig. 1b). This may reflect a selective pressure for mice with lower levels of Parp1 activity for embryonic viability, because viable *Xrcc1^Nes-cre^* mice are born at a sub-mendelian ratio *(17)*.

**Figure 1.**
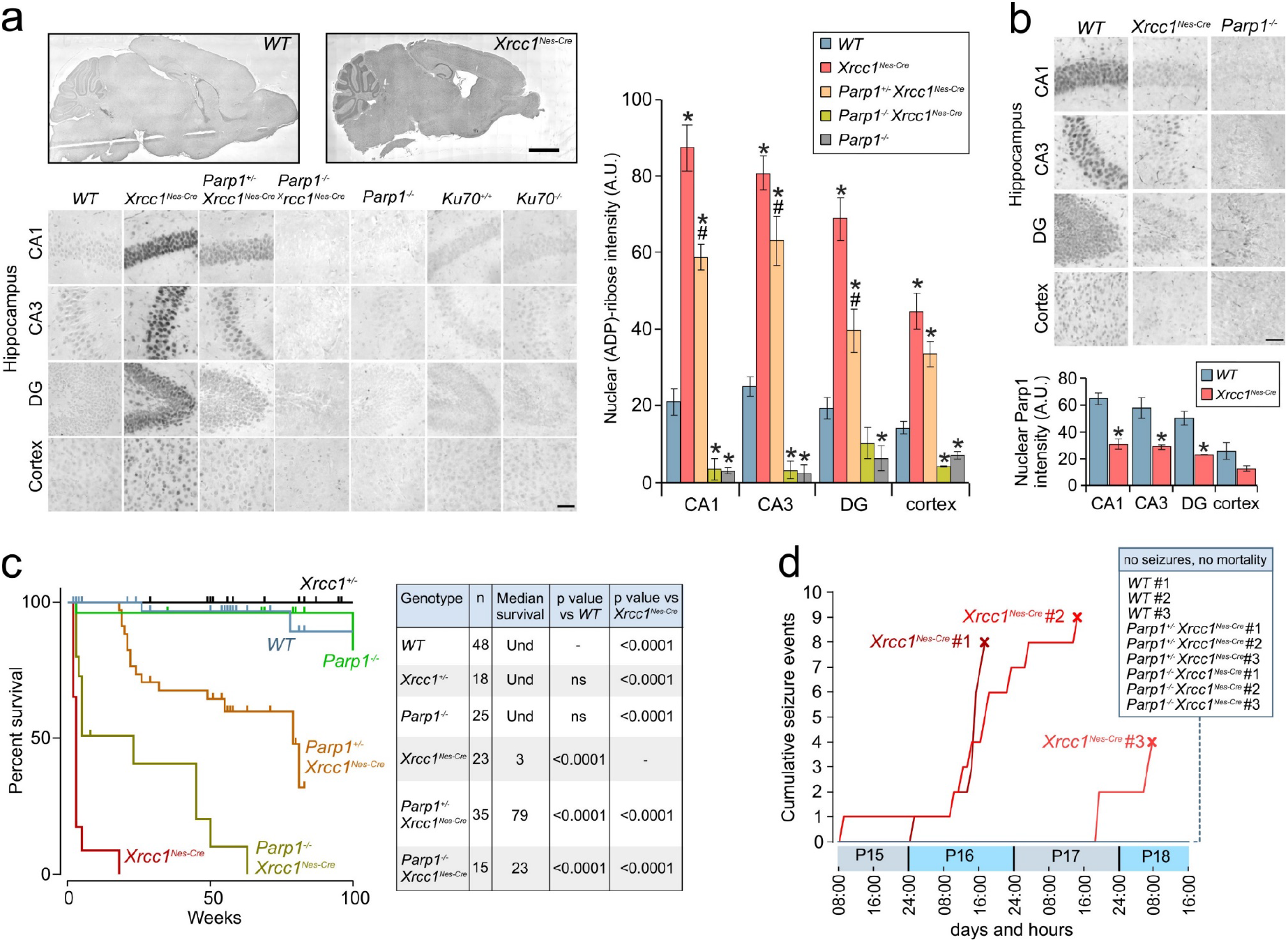
Parp1 hyperactivation triggers juvenile seizures and mortality in the absence of Xrcc1. **[A]** Sagittal sections obtained from mice of the indicated genotypes were immunostained for ADP-ribose. Representative images of sagittal sections showing levels of ADP-ribose in the hippocampal regions CA1, CA3, and dentate gyrus (DG), and in the cerebral cortex. Scale bars: 5 mm, 50 μm. *WT*(n = 4 mice), *Xrcc1^Nes-Cre^* (n = 4), *Parp1^+/-^/Xrcc1^Nes-Cre^* (n = 4), *Parp1^-/-^/Xrcc1^Nes-Cre^* (n = 3) and *Parp1^-/-^* (n = 3). Summary histograms show mean ± SEM. ANOVA single factor between genotypes in CA1: p = 2.2×10^-10^, CA3: p = 1.7×10^-9^, DG: p = 2.8×10^-7^, cortex: p = 2.4×10^-6^. * and # indicate t-test comparisons where p < 0.05. * is for comparisons of *WT* versus other genotypes. # is for comparisons of *Xrcc1^Nes-Cre^* versus *Parp1^+/-^ /Xrcc1^Nes-Cre^*. **[B]** Representative images of Parp1 based on immunostaining in hippocampus and cortex. Bottom panels show quantification (mean ± SEM). The level of Parp1 is significantly reduced in *Xrcc1^Nes-Cre^* brain versus wild type (n = 3 mice per genotype. t-test * p < 0.02). **[C]** Kaplan-Meier curve for survival of mice of the indicated genotypes. The table shows number of individuals in each group, median survival in weeks, and p-values from pairwise curve comparisons; Und: undetermined (Log-rank Mantel-Cox tests). **[D]** Video monitoring and recording of generalised running/bouncing seizures in mice of the indicated genotypes from P15-P19. The point of death of *Xrcc1^Nes-Cre^* mice by fatal seizure is indicated (cross). Note no seizures from mice of other genotypes were detected. *WT* (n = 9 mice), *Xrcc1^Nes-cre^* (n = 3 mice), *Parp1^+/-^/Xrcc1^Nes-Cre^* (n = 3 mice), and *Parp1^-/-^/Xrcc1^Nes-Cre^* (n = 3).

*Xrcc1^Nes-Cre^* mice exhibit greatly reduced longevity, with the cohort employed in these experiments having a median lifespan of ~3-4 weeks (Fig.1C). Given the widespread PARP1 hyperactivity across the brain in *Xrcc1^Nes-Cre^* mice, we examined the impact of Parp1 deletion on longevity. Strikingly, the median lifespan of *Parp1^+/-^ /Xrcc1^Nes-Cre^* and *Parp^-/-^/Xrcc1^Nes-Cre^* littermates in which one or both *Parp1* alleles were additionally deleted was increased ~7-25 fold; to 23 and 79 weeks, respectively (Fig.1C). This result is significant because it indicates that aberrant Parp1 activity in Xrcc1-defective brain is a major contributor to organismal death. Interestingly, deletion of one *Parp1* allele prolonged the lifespan of *Xrcc1^Nes-Cre^* mice to a greater extent than deletion of both *Parp1* alleles. This result suggests that whilst the loss of a single *Parp1* allele is sufficient to suppress Parp1-induced toxicity, the loss of the second allele eradicates an additional role for Parp1 in Xrcc1-defective brain that is important for survival.

To identify the cause of Parp1-dependent death in *Xrcc1^Nes-Cre^* mice we conducted infrared video imaging for a four-day period starting at P15. These experiments revealed that *Xrcc1^Nes-Cre^* mice experience sporadic seizures, culminating ultimately in a lethal seizure from which they did not recover (Fig.1D). In contrast, we did not observe any seizures in wild type mice over the same time period. The cause of death induced by the seizures is unclear, but similar to Sudden Unexpected Death During Epilepsy (SUDEP) in humans it is likely to result from the disruption of normal cardiac or respiratory function *(18)*. Importantly, we also did not detect seizures in *Xrcc1^Nes-Cre^* mice in which one or both alleles of Parp1 were deleted, over the time course of the experiment, consistent with greatly elevated lifespan of these mice (Fig.1D). To our knowledge, this is the first demonstration that seizures can be triggered by aberrant Parp1 activity.

The presence of high levels of poly(ADP-ribose) in *Xrcc1^Nes-Cre^* hippocampus was of particular interest, because defects in this region of the brain are often associated with seizure activity *(19)*. To examine directly whether *Xrcc1^Nes-Cre^* hippocampus exhibits elevated seizure-like activity we conducted electrophysiological experiments in acute brain slices using a high-density multi-electrode array platform that could assay network activity simultaneously across hippocampal and cortical structures (Fig. 2A). Continuous recordings were made as slices were perfused into an epileptogenic solution. We found that cumulative seizure-like activity was indeed significantly higher in *Xrcc1^Nes-Cre^* hippocampus versus wild type, with the strongest effects observed in CA3 region (Fig.2B-D). To examine this further, we carried out targeted extracellular recording experiments in CA3 and tested the effect of *Parp1* deletion on seizure expression. Again, after washing into an epileptogenic solution, the mean cumulative number of seizure-like events was ~2.5-fold greater in *Xrcc1^Nes-Cre^* than in wild type mice (Fig.2E-G). Moreover, this elevated seizure-like activity was reduced and/or prevented if one or both alleles of *Parp1* were deleted, implicating Parp1 hyperactivity as a cause of the elevated seizure activity in *Xrcc1^-/-^* hippocampus, *in vitro* (Fig.2E-G).

**Figure 2.**
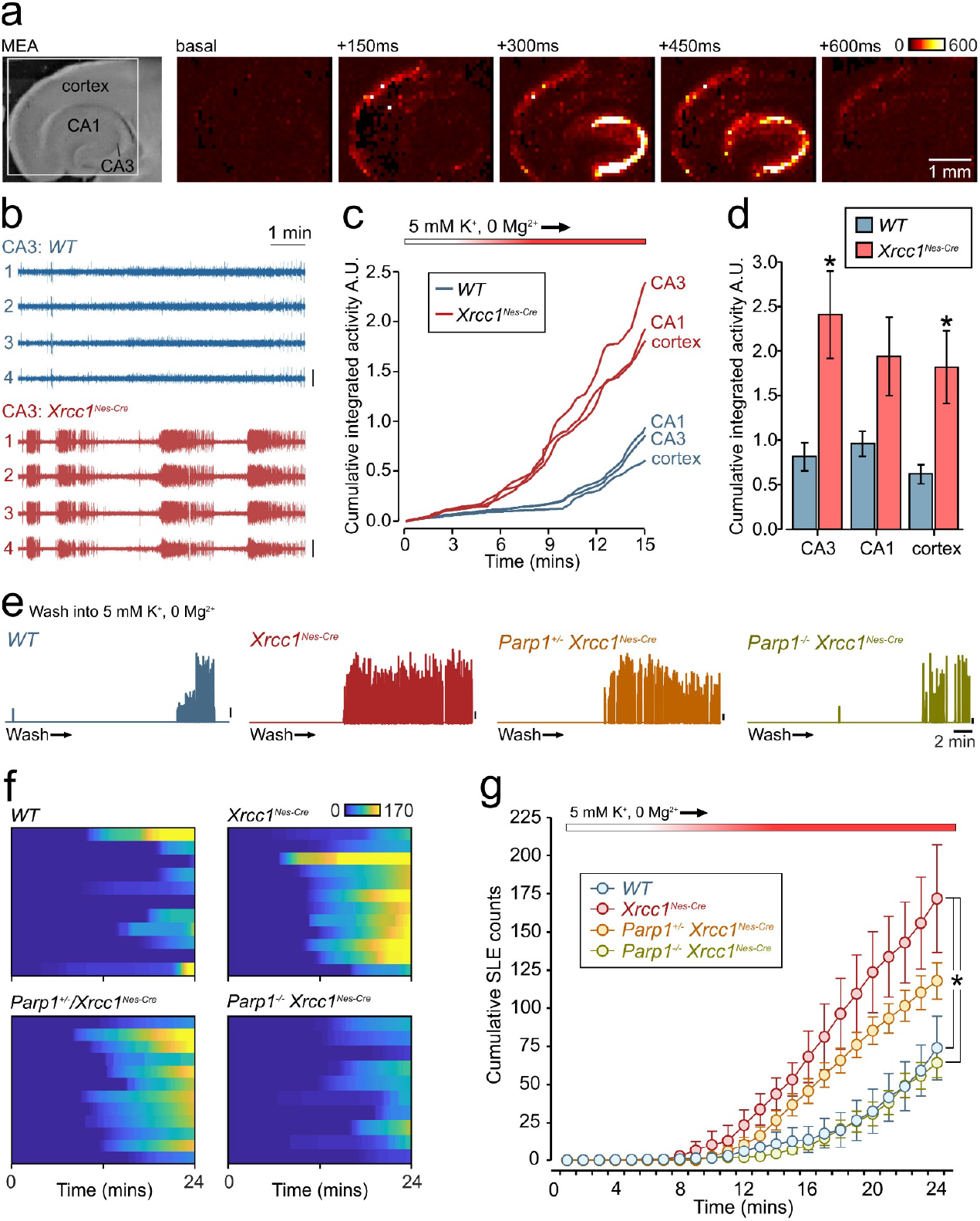
Low seizure threshold in *Xrcc1^Nes-Cre^* brain slices is corrected by additional *Parp1* deletion. **[A]** Brightfield image (left) shows acute brain slice positioned on HD-MEA with target hippocampal and cortical regions indicated. (Right) progression of a typical seizure event when slice is perfused with an epileptogenic solution; colour-code indicates voltage changes in microvolts. [**B**] Representative traces of four channels in CA3 region of hippocampus at 10 min timepoint with seizure-like activity in *Xrcc1^Nes-Cre^* but not *WT*. **[C]** Mean cumulative activity plots in CA1 and CA3 regions of the hippocampus and the cortex over 15 minutes of recording during perfusion into epileptogenic buffer. *WT* (n = 8 slices), *Xrcc1^Nes-Cre^* (n = 9). **[D]** Summary histogram shows mean ± SEM for cumulative activity at 15 min timepoint. CA3 and cortex have significantly higher activity in the *Xrcc1^Nes-Cre^* brain slices compared to control (* p < 0.05, unpaired t-tests). **[E]** Plots show enveloped waveforms of targeted extracellular recordings from CA3 region during perfusion into epileptogenic buffer for *WT, Xrcc1^Nes-Cre^*, *Parp1^+/-^/Xrcc1^Nes-Cre^* and *Parp1^-/-^/Xrcc1^Nes-Cre^*. Upward deflections indicate seizurelike episodes (SLEs). Scale bars, 5% of maximum amplitude. [**F**] Heatplot summaries of cumulative SLE counts for all conditions: *WT* (n = 11 slices), *Xrcc1^Nes-Cre^* (n = 12), *Parp1^+/-^ /Xrcc1^Nes-Cre^* (n = 12) and *Parp1^-/-^/Xrcc1^Nes-Cre^* (n = 10). Each horizontal bar corresponds to one hippocampal slice with colour-code indicating cumulative SLEs. **[G]** Summary of mean ± SEM cumulative SLE counts for each condition. At the recording endpoint, the total SLE count in *Xrcc1^Nes-Cre^* is significantly higher than both *WT* and *Parp1^-/-^ Xrcc1^Nes-Cre^* (Kruskal-Wallis ANOVA, p < 0.0017, * indicates significance with Dunn’s post-hoc comparisons).

To address the mechanistic basis of the seizure-like activity triggered by Parp1 we isolated hippocampal neurons from *Xrcc1^Nes-Cre^* mice. Similar to brain sections, we detected elevated endogenous levels of ADP-ribosylation in *Xrcc1^Nes-Cre^* neurons, although in these isolated cells detection of this signal required incubation with an inhibitor of poly(ADP-ribose) glycohydrolase (PARG); the enzyme primarily responsible for poly(ADP-ribose) catabolism (Fig.3A & 3B). To confirm that the poly(ADP-ribose) detected here was nascent polymer synthesised during incubation with PARGi, rather than pre-existing poly(ADP-ribose), we co-incubated the neurons with an inhibitor of PARP1 (PARPi). Indeed, PARPi ablated the appearance of poly(ADP-ribose) in the *Xrcc1^Nes-Cre^* neurons (Fig.3A & 3B). This result indicates that Parp1 hyperactivation occurs continuously in *Xrcc1^Nes-Cre^* hippocampal neurons, presumably as a result of the elevated steady-state level of unrepaired SSBs.

**Figure 3.**
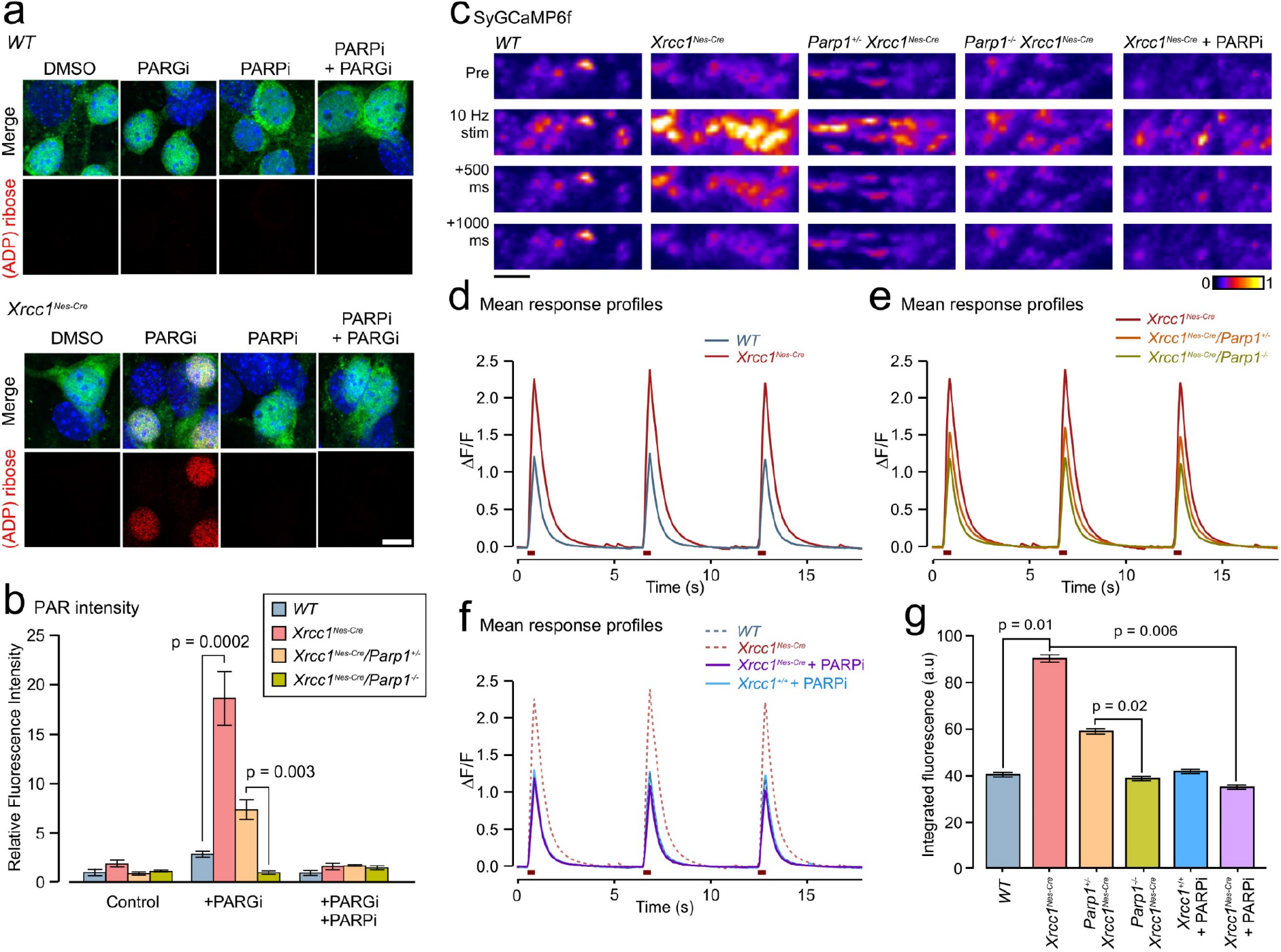
Aberrant presynaptic calcium signaling in *Xrccl^-/-^* hippocampal neurons is rescued by Parp1 deletion or inhibition. **[A]** Indirect immunofluorescence of DIV6 hippocampal neurons cultured from P1 *WT* and *Xrcc1*^Nes-Cre^ mouse pups. Cells were pretreated with PARP inhibitor (10 μM) or vehicle for 2 hr prior to fixation, with PARG inhibitor (10 mM) additionally present for the final hour. Scale bar 10 μm. **[B]** Histogram of mean relative PAN fluorescence (quantified manually in ImageJ) in MAP2-positive hippocampal neurons pretreated with PARP inhibitor (5 μM) or vehicle for 5 h prior to fixation, with PARG inhibitor (10 mM) additionally present for the final hour. Neurons were cultured from *WT* (n = 6 mice, > 180 cells per condition), *Xrcc1^Nes-Cre^* (n = 6, > 180), *Parp1*^+/-^/*Xrcc1*^Nes-Cre^ (n = 3, > 90), and *Parp1^-/-^/Xrcc1^Nes-Cre^* (n = 3, > 90). * indicates significant differences from *WT* (Kruskal-Wallis ANOVA, p = 0.0013 and Dunn’s post-hoc tests) **[C]** Representative images of fluorescence responses in synaptic terminals expressing SyGCaMP6f to 10 AP (10 Hz) stimulation in dissociated hippocampal neurons derived from wild type and mutant mice. Scale bar: 5 μm. **[D,E]** Mean SyGCaMP6f responses to three rounds of 10 APs stimulation from mice of the following genotypes; *WT* (n = 1946 synapses, 9 coverslips, 3 animals), *Xrcc1^Nes-Cre^* (n = 3313, 12, 4), *Parp1^+/-^ /Xrcc1^Nes-Cre^* (n = 2272, 9, 3), and *Parp1^-/-^/Xrcc1^Nes-Cre^* (n = 2122, 11, 3). **[F]** Response to chronic treatment with the PARP inhibitor, KU 0058948 Hydrochloride (1 μM) for 9-11 days prior to imaging for *Xrcc1^+/+^* (n = 1264, 8, 3 and *Xrcc1^Nes-Cre^* (n = 2231, 10, 4) mice. Dashed lines show *WT and Xrcc1^Nes-Cre^* for comparison. **[G]** Summary histogram of mean ± SEM of total integrated fluorescence responses to stimulation for each condition. Responses are significantly higher in *Xrcc1^Nes-Cre^* synapses versus both *WT* and *Parp1^-/-^/Xrcc1^Nes-Cre^* (oneway ANOVA, p < 0.0033, and pairwise student’s t-tests)

Currently, nothing is known about the mechanisms by which aberrant Parp1 activity at unrepaired SSBs can trigger seizures. However, it is well established that Parp1 activity reduces intracellular levels of its co-factor NAD+, which in turn can affect the expression of many genes including those affecting Ca^2+^ homeostasis and neurodegeneration *(20)*. Consequently, we examined whether the seizure-like activity in *Xrcc1^Nes-Cre^* mice reflects a defect in Ca^2+^ signalling, at the level of single synapses. To test this idea, we expressed SyGCaMP6f, a presynaptically-targeted optical Ca^2+^ reporter, in dissociated hippocampal cultures from different genotypes and imaged fluorescence changes in response to electrical stimulation. With repeated presentations of 10 Hz stimulus trains, SyGCaMP6f-positive puncta in wild type mouse cultures showed characteristic transient increases in fluorescence consistent with activity-evoked Ca^2+^ influx at the presynaptic terminal (Fig.3C). Strikingly, however, the amplitude of these responses was ~2-fold higher in *Xrcc1^Nes-Cre^* neurons and was reduced or suppressed to wild type levels if one or both alleles of *Parp1* were deleted (Fig.4C-G). This demonstrates that Parp1 hyperactivity results in excessive activity-evoked synaptic Ca^2+^ influx. To our knowledge, this is the first demonstration that aberrant Parp1 activity deregulates synaptic Ca^2+^ signalling. To confirm this idea, we incubated cultures with a PARP1 inhibitor (PARPi) starting 9-11 days prior to recording. Strikingly, this treatment fully suppressed the aberrant Ca^2+^ response in *Xrcc1^Nes-Cre^* neurons back to wild type level (Fig.4F-G). Collectively, our results demonstrate that aberrant and/or excessive Parp1 activity triggers elevated Ca^2+^ signalling at presynaptic terminals in response to unrepaired SSBs, providing a compelling explanation for the elevated seizures and, consequently, shortened lifespan in *Xrcc1^Nes-Cre^* mice.

In summary, we reveal here that Parp1 hyperactivity is a major molecular mechanism by which seizures and shortened lifespan are triggered in *Xrcc1^Nes-Cre^* fmice. We show that aberrant Parp1 activity triggers seizure-like activity in Xrcc1-defective hippocampus *ex vivo* and excessive presynaptic Ca^2+^ signalling in hippocampal neurons *in vitro*; both of which are supressed by Parp1 deletion or inhibition. These data link aberrant Parp1 activity at DNA single-strand breaks to deregulated calcium homeostasis and lethal seizures and highlight Parp inhibitors as a possible approach for the treatment of XRCC1-defective disease.

## Acknowledgements

We thank Zhao-Qi Wang for the *Parp1^-/-^* mouse strain and Tom Baden and Marvin Seifert for their support with the MEA work. **Funding:** This work was funded by an MRC Programme grant to KWC and KS (MR/P010121/1) and a BBSRC Project grant to KS (BB/K019015/1). KWC is the recipient of a Royal Society Wolfson Research Merit Award. **Competing interests**: Authors declare no competing interests. **Data and materials availability**: All data is available in the main text and materials are available on request.

## Materials and Methods

### Animals and animal care

Experiments were carried out in accordance with the UK-Animal (Scientific Procedures) Act 1986 and satisfied local institutional regulations at the University of Sussex. Mice were maintained and used under the auspices of UK Home Office project license number P3CDBCBA8. The generation of *Parp1^-/-^, Xrcc1^Nes-Cre^*, and *Ku70^-/-^* mice were reported previously *(17,21)*. Intercrosses between *Parp1^-/-^* and *Xrcc1^+/loxp^* mice were maintained in a mixed background C57Bl/6 × S129 strain and housed on a 12h light/dark cycle with lights on at 07:00. Temperature and humidity were maintained at 21 °C (± 2°C) and 50% (± 10%), respectively. All experiments were performed under the UK Animal (Experimental Procedures) Act, 1986.

### Immunohistochemistry and microscopy

Antibodies used were rabbit Fc-fused Anti-pan-ADP-ribose binding reagent Millipore; MABE1016), anti-PARP1 mouse monoclonal (Serotec; MCA1522G), anti-NeuN mouse monoclonal (Millipore MAB337), and anti-MAP2 chicken (Abcam; ab5392). Secondary antibodies used for immunofluorescence were goat anti-rabbit Alexa 647, anti-mouse Alexa 488 (Invitrogen; A21244 and A11001), and anti-chicken Alexa 488 (Abcam; ab150173) and, for immunohistochemistry, Biotin-SP-conjugated AffiniPure goat anti-rabbit antibody (JacksonImmuno Research; 111-065-144). Mice were anaesthetized using 0.25 mg/g Dolethal (Vetoquinol UK Ltd) and perfused transcardially with PBS followed by 4% formaldehyde. Brains were postfixed in 4% paraformaldehyde for 48 h and stored in 25% sucrose/PBS until moulding and freezing (TFM-5). 10-20 μm sagittal sections were prepared using a cryostat (Leica CM1850). Immunohistochemistry was conducted as described previously *(17)*. Fluorescent images were acquired using the Zeiss LSM880 confocal microscope, with oil immersion objective (Plan-Apochromat 63×/1.4 Oil DIC M27). The Airyscan super-resolution module was used to obtain high-resolution images. The image stacks were processed for brightness and contrast in ImageJ 1.51j. Images of immunohistochemistry samples were acquired with LSM880 using the wide-field imaging mode (Axiocam 503 Mono, Plan-Apochromat 10x/0.45 and 20x/0.8 M27). Nuclear staining was quantified manually using ImageJ 1.51j. The mean nuclear signal was normalized by subtracting the mean value for background staining. Whole tissue sections were imaged in a tiling mode with 10% overlap and stitched in Zeiss Zen.

### Electrophysiology

For targeted extracellular recordings, acute transverse hippocampal slices (300 μm) were prepared from P14-P16 mice using a vibroslicer (VT1200S, Leica Microsystems, Germany) in ice-cold artificial cerebrospinal fluid containing (in mM): 125 NaCl, 2.5 KCl, 25 glucose, 1.25 NaH_2_PO_4_, 26 NaHCO_3_, 1 MgCl_2_, 2 CaCl_2_ (bubbled with 95% O_2_ and 5% CO_2_, pH 7.3). All experiments were performed at 23-25·C. During an experiment, an extracellular electrode was placed in the hippocampal CA3 pyramidal region and field voltage recordings made as slices were perfused from ACSF into a modified (epileptogenic) saline containing (in mM) 125 NaCl, 5 KCl, 25 glucose, 1.25 NaH_2_PO_4_, 26 NaHCO_3_, 2 CaCl_2_. Signals were amplified using a MultiClamp 700A (Molecular Devices), digitized at 50 kHz with a Digidata 1320 and recorded in pCLAMP acquisition software (Molecular Devices) for offline analysis. To quantify seizure-like activity, raw traces were exported and analyzed in Matlab (Mathworks). Root mean square signal envelopes (sliding window length: 400 samples) were calculated for each trace and a peak waveform generated using an automated peak-find search function. This waveform was then enveloped (sliding window length: 1500 samples) and a peak count analysis used to quantify seizure-like episodes (SLEs) for each sample. To independently verify our automated analysis approach, we also carried out a separate manual count of SLEs, by tallying episodes in 2.5 s time bins. Outputs from both types of quantification were highly significantly positively correlated (Spearman-rank, *r_s_* = 0.833, p < 0.0001). For MEA recordings, slices were placed onto a high-density Stimulo MEA chip (4096 electrodes: size 21 x 21 μm, pitch 81 μm, 64 x 64 matrix, 3Brain). The slices were immobilized using a custom-made weight under membrane and were constantly perfused with oxygenated (epileptogenic) ACSF (as above) at +34°C. Recordings were acquired with BrainWave v4.2 software in 10 min time windows, digitized at 9.5 kHz and stored for offline analysis. 7-channel arrays from target regions were batch exported in 3brain HDF5 format and quantified in Matlab. Signals were processed first by enveloping (RMS signal envelope, sliding window length: 500 samples) and then by generating a peak waveform to identify seizure-like events. These waveforms were then integrated to provide a collective measure of burst frequency and amplitude at each timepoint and presented as cumulative totals.

### Videoanalysis

Videomonitoring was performed using the Noldus PhenoTyper 3000 system, including infrared LED units and a video recording camera for the duration of the experiment (Video output CCIR black/white VPP −75 Ohm (PAL) or EIA black/white Vpp-75 Ohm (NTSC)). Mice were placed in the chamber with floor area 30cm x 30cm and were provided with bedding, minimal nesting, food pellets and a water source. Mouse pups of the indicated genotype were housed with mother and a control sibling from P15 up to P20. Video recordings were observed after recording and the number of running-bouncing-seizures was quantified.

### Cell Culture

P1-2 mouse pups were decapitated, brains were removed and placed in ice-cold Hank’s Buffered Saline Solution (HBSS), 0.1M HEPES. Hippocampi were isolated and then washed three times in warmed 10% FCS, 20 mM Glucose, 1% Pen/Strep Minimal Essential Media (MEM) (Gibco). Hippocampi were manually dissociated by trituration and diluted to a density of 50,000 cells/well and plated on poly-D-Lysine (20μg/ml) coated 15 mm glass coverslips. 2 h after plating, the media was replaced with 2% B27, 1% Glutamax, 1% Pen/Strep-supplemented Neurobasal A medium (Gibco). Cells were maintained without cytosine arabinoside, typically used to manage glial cell count, to prevent induction of exogenous DNA damage (22). Cells were maintained in 5% CO_2_ at 37°C and fed every 3 d through half-exchange of media. Hippocampal cells were taken for ICC experiments at DIV6, and live imaging at DIV15-17. Hippocampal cells treated for PAN immunofluorescence were administered 10 μM PARG inhibitor PDD 00017273 (Tocris) in DMSO for 1 h following pre-treatment with 5 μM PARPi Ku-0058948 or DMSO vehicle. Cells were fixed in 4% paraformaldehyde and permeabilized with 0.2% Triton-X for 2 min. Cells were blocked in 10% Goat Serum, 0.3% Triton-X for 30 minutes prior to incubation with relevant primary antibodies. Cells were incubated with relevant secondary antibodies following PBST washes, and counterstained with DAPI prior to mounting.

### SyGCaMP6f Imaging

Hippocampal neurons were transduced with AAV6_SyGCaMP6f at DIV6/7 at a multiplicity of infection (MOI) of 100. Images were acquired using a 60x/1.0 NA objective on an Olympus BX61WI microscope fitted with an Andor Ixon+ EM-CCD (40 ms exposure, 20 Hz acquisition frequency and 4×4 binning) controlled by custom-written Micromanager routines. Coverslips were placed in a custom-built imaging chamber and a Grass SD9 Stimulator was used to apply field stimulation (voltage: 22.5V, 1 ms pulse width). All experiments were carried out in Extracellular Bath Solution (EBS) containing (in mM), 136 NaCl, 10 HEPES, 10 D-Glucose, 2.5 KCl, 2 CaCl_2_, 1.3 MgCl_2_. EBS was supplemented with 50 μM APV and 20 μM CNQX to inhibit NMDA and AMPA receptors respectively. Image stacks were analyzed using IGOR Pro 8. Stacks were imported via the SARFIA plugin (23) and corrected for x-y drift using the built-in image registration function (24). Regions of Interest (ROIs) were detected via image segmentation using a threshold of (−3) x SD of all pixel values in the Laplace operator. Mean background intensity was subtracted to account for alterations to background fluorescence and ΔF/F values were calculated using the 10 frames prior to stimulation as a baseline. A custom script was used to determine whether ROIs had exceeded a threshold change in fluorescence intensity following stimulation and ROIs that failed to reach threshold intensity were removed from the mask.

